# Broad neutralization against SARS-CoV-2 variants induced by a modified B.1.351 protein-based COVID-19 vaccine candidate

**DOI:** 10.1101/2021.05.16.444369

**Authors:** Danmei Su, Xinglin Li, Cui He, Xueqing Huang, Meilin Chen, Qiang Wang, Wenchang Qin, Ying Liang, Rong Xu, Jinhua Wu, Peiwen Luo, Xiaofang Yang, Yilan Zeng, Mei Luo, Dongxia Luo, David M Salisbury, Donna Ambrosino, George Siber, Ralf Clemens, Peng Liang, Joshua G. Liang

## Abstract

Beginning in late 2020, the emergence and spread of multiple variant SARS-CoV-2 strains harboring mutations which may enable immune escape necessitates the rapid evaluation of second generation COVID-19 vaccines, with the goal of inducing optimized immune responses that are broadly protective. Here we demonstrate in a mouse immunogenicity study that two doses of a modified B.1.351 spike (S)-Trimer vaccine (B.1.351 S-Trimer) candidate can induce strong humoral immune responses that can broadly neutralize both the original SARS-CoV-2 strain (Wuhan-Hu-1) and Variants of Concern (VOCs), including the UK variant (B.1.1.7), South African variant (B.1.351) and Brazil variant (P.1). Furthermore, while immunization with two doses (prime-boost) of Prototype S-Trimer vaccine (based on the original SARS-CoV-2 strain) induced lower levels of cross-reactive neutralization against the B.1.351 variant, a third dose (booster) administered with either Prototype S-Trimer or B.1.351 S-Trimer was able to increase neutralizing antibody titers against B.1.351 to levels comparable to neutralizing antibody titers against the original strain elicited by two doses of Prototype S-Trimer.

## Introduction

Despite the tremendous progress and unprecedented speed of development of COVID-19 vaccines, a new record for daily COVID-19 cases was set in April 2021 (*1*), over 16 months after the SARS-CoV-2 virus outbreak first emerged. Total deaths caused by COVID-19 surpassed 3 million in May 2021, with 1 million deaths having accumulated in the prior 3 months alone.

Most concerning is the emergence of multiple new SARS-CoV-2 variants of concern (VOC) starting in late 2020 that is concurrent with a surge in COVID-19 cases globally. These VOCs appear to be associated with mutations in the spike (S) protein which could potentially increase the rate of viral transmission and/or escape immunity induced by vaccination with first-wave COVID-19 vaccines based on the original strain of SARS-CoV-2 (Wuhan-Hu-1) which was first published in January 2020 (*2, 3*). The emergence and spread of the B.1.1.7 variant in the United Kingdom (UK), B.1.351 variant in South Africa and P.1 variant in Brazil have led to their classification as VOCs. These VOCs all include the N501Y mutation in the receptor-binding domain (RBD) of the S protein that is reported to increase transmission by 40% to 70% (*2*). The B.1.351 and P.1 variants have two additional RBD mutations – E484K and K417 – that may allow immune escape from antibodies induced by prototype vaccines and natural infection (*3*).

Preliminary evidence from randomized, controlled clinical trials of prototype COVID-19 vaccines suggest decreased vaccine efficacy against VOCs compared to original SARS-CoV-2 strains. NVX-CoV2373, an adjuvanted protein-based COVID-19 vaccine, demonstrated 89% efficacy in the UK (where B.1.1.7 predominates) but only 49% efficacy in South Africa (where B.1.351 predominates) (*4*). ChAdOx1, an adenovirus-vectored COVID-19 vaccine, demonstrated only 10% efficacy against the B.1.351 variant (*5*). BNT162b2, an mRNA-based COVID-19 vaccine, demonstrated 95% efficacy against COVID-19 caused predominantly by the original SARS-CoV-2 strains (*6*), but demonstrated about 75% efficacy against the B.1.351 variant (*7*). These observations of decreased vaccine efficacy against B.1.351 correspond with significantly decreased neutralizing antibody titers against the B.1.351 variant compared to the original strain in vaccinee sera (*5, 8, 9,10*). Subjects vaccinated with CoronaVac, an inactivated SARS-CoV-2 vaccine, demonstrated no detectable neutralizing antibody titers against the P.1 variant (*11*).

Lower vaccine efficacy against VOCs combined with their increased transmission rates could make achieving herd immunity particularly difficult, a problem that could be further exacerbated by the shortage of COVID-19 vaccines globally, especially in low- and middle-income countries (*12,13*). If not effectively controlled, the rapid global spread of SARS-CoV-2 VOCs could potentially lead to the continued emergence of new variants of interest or VOCs harboring new escape mutations, such as the Indian variant (B.1.617) which emerged concurrent with a massive spike in COVID-19 cases in the spring of 2021 and has now been declared a new VOC by the WHO (*14,15*).

These circumstances necessitate the rapid evaluation of booster dose strategies in order to enhance neutralizing antibody responses to VOCs and the development of second generation COVID-19 vaccines which may confer optimized broadly-protective, cross-neutralizing characteristics. In this study, we produced a modified B.1.351 Spike -Trimer antigen (dubbed “B.1.351 S-Trimer”) and compared it with our Prototype S-Trimer (*16,17*) in a mouse immunogenicity study. Heterologous prime-boost and bivalent vaccine approaches were also evaluated, as well as the effect of a third dose (booster dose).

## Results

### Detection of cross-reactive neutralizing antibodies in human COVID-19 convalescent sera

8 human convalescent serum (HCS) samples with moderate-to-high antibody titers collected from COVID-19 patients infected with the original strain (table s1) were analyzed for SARS-CoV-2 pseudovirus neutralization titers (Fig. 1A) and ACE2-competitive ELISA titers (Fig. 1B) against the original strain, UK variant (B.1.1.7), South African variant (B.1.351) and Brazil variant (P.1) strains. Neutralizing antibody titers in HCS against B.1.1.7 appeared to be similar to the original strain, and differences in ACE2-competitive titers were also not statistically significant. However, significantly lower antibody titers in HCS were observed against B.1.351 (7- to 9-fold lower) and P.1 (2- to 3-fold lower) compared to titers against the original strain. These results appear to be consistent with published data (*3,9*), supporting the use of these assays in further animal vaccination studies.

**Fig. 1.**
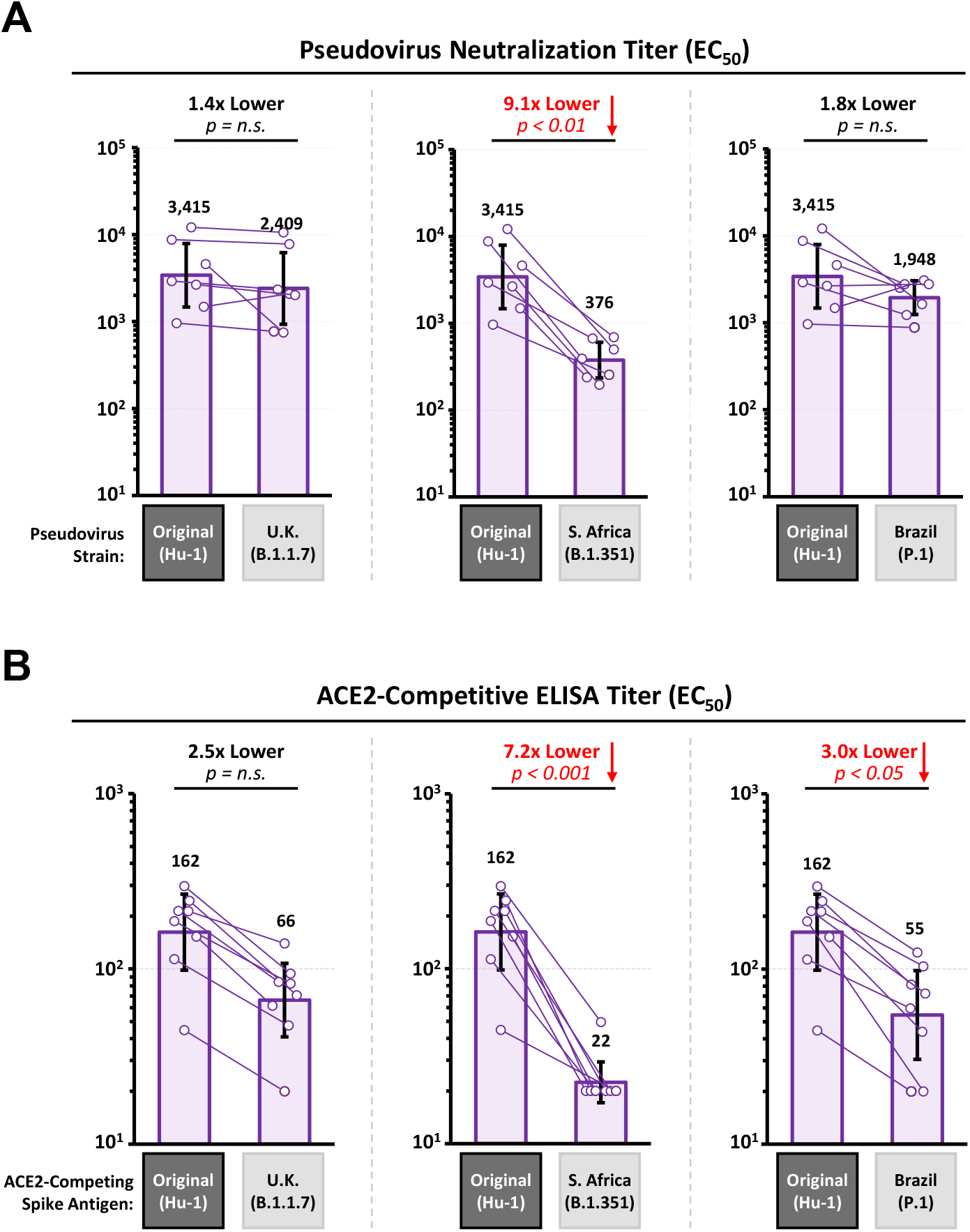
Detection of SARS-CoV-2 Neutralizing Antibodies and ACE2-Competitive Titers in Human COVID-19 Convalescent Sera. 8 human convalescent sera (HCS) samples with moderate-to-high antibody titers collected from COVID-19 patients infected with original strain (Wuhan-Hu-1) were analyzed for (**a**) SARS-CoV-2 pseudovirus neutralization titers and (**b**) ACE2-competitive ELISA titers based on the original (Wuhan-Hu-1), UK (B.1.1.7), South African (B.1.351) and Brazil (P.1) strains. Data are presented comparing variant strains titers to original strain titers. Dots represent data for individual HCS samples, and bars represent geometric mean titers (GMT) of half maximal effective concentration (EC_50_) values. Error bars represent 95% confidence intervals (95% CI).

### Production and characterization of spike antigens based on original and B.1.351 strains

Covalently-trimerized S-protein antigens were produced based on the original SARS-CoV-2 strain (Prototype S-Trimer) and South African variant (B.1.351 S-Trimer) utilizing Trimer-Tag^©^ technology (*16*). Prototype S-Trimer is based on the native spike ectodomain sequences of the original SARS-CoV-2 strain (Wuhan-Hu-1), whereas the modified B.1.351 S-Trimer contains 3 RBD mutations (K417N, E484K, N501Y) and the D614G mutations found in the B.1.351 variant strain, while the N-terminal domain (NTD) and S2 sequences are based on the original strain (Fig. 1A).

The binding affinity (K_D_) of purified S-Trimer antigens to the human ACE2 receptor using ForteBio BioLayer interferometry was shown to be approximately 5.2 nM for Prototype S-Trimer and 1.5 nM for B.1.351 S-Trimer (Fig. 1B). The 3- to 4-fold higher ACE2 binding affinity of the B.1.351 S-Trimer compared to Prototype S-Trimer is similar to previously-reported results (*18,19,20*), which appears to be due mainly to a slower off-rate (K_off_) and is speculated to further exacerbate the lower potency observed in B.1.351 cross-neutralization assays, since antibodies of lower affinity may struggle to compete with B.1.351 spike protein for ACE2 binding.

### Immunogenicity of Prototype and B.1.351 S-Trimer prime-boost (2 dose) regimens in mice

BALB/c mice were immunized in Stage 1 of the study (Fig. 2C, 2D) with either two doses (Day 0 and Day 21) of Prototype S-Trimer, heterologous prime-boost (Dose 1 Prototype S-Trimer + Dose 2 B.1.351 S-Trimer), two doses of B.1.351 S-Trimer, or two doses of bivalent vaccine (Prototype S-Trimer mixed with B.1.351 S-Trimer). All animals in Stage 1 received a priming dose (Dose 1) adjuvanted with CpG 1018 plus alum. For the second dose (boost), half of the animals in each group received a boost adjuvanted with CpG 1018 plus Alum, and the other half received non-adjuvanted boost (antigen-only). Humoral immunogenicity analysis was conducted on Day 35 blood samples (2 weeks post-dose 2) based on pseudovirus neutralization titers against the original strain and three VOCs (B.1.1.7, B.1.351 and P.1).

**Fig. 2.**
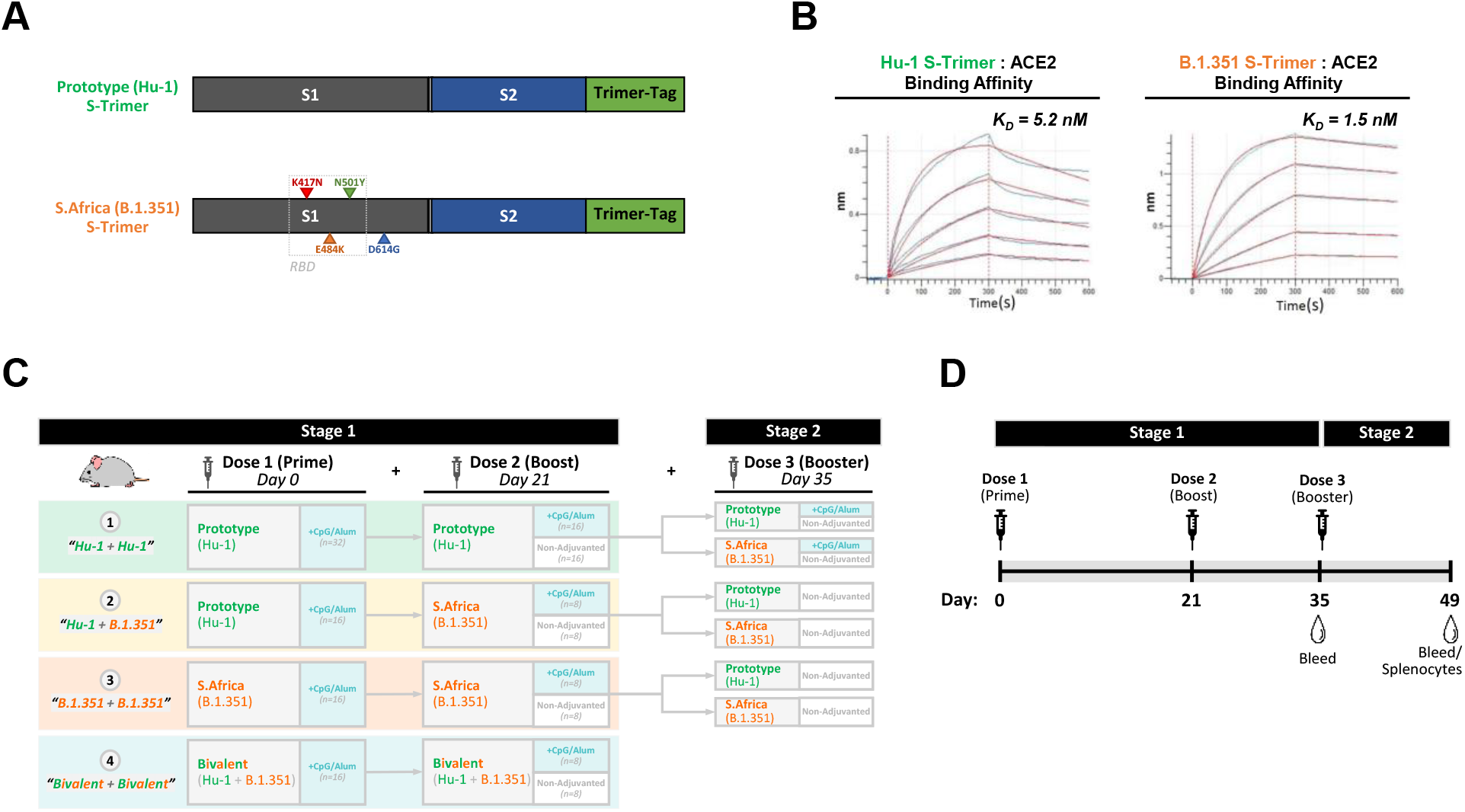
Prototype and Modified B.1.351 S-Trimer Antigens and Mouse Immunogenicity Study Design. (**a**) Schematic representations of Prototype S-Trimer and the modified B.1.351 S-Trimer containing 3 RBD mutations and the D614G mutations in the B.1.351 variant. (**b**) Determination of the binding affinity (K_D_) between S-Trimers (Prototype and B.1.351) and human ACE2-Fc by ForteBio BioLayer interferometry. (**c**) BALB/c mice (n=16-32/group) were immunized in Stage 1 of the study with either two doses of Prototype S-Trimer (3 µg), heterologous prime-boost (dose 1 Prototype S-Trimer; dose 2 B.1.351 S-Trimer; 3 µg of each antigen), two doses B.1.351 S-Trimer (3 µg), or two doses of bivalent vaccine (3 µg Prototype S-Trimer mixed with 3 µg B.1.351 S-Trimer). All animals in Stage 1 received a priming dose (Dose 1) adjuvanted with CpG 1018 plus alum. For the boost (Dose 2), half of the animals in each group received a boost adjuvanted with CpG 1018 plus Alum, and the other half received non-adjuvanted boost (antigen-only). In Stage 2 of the study, animals in group 1 were randomized to receive a booster dose (Dose 3) with either 3 µg Prototype S-Trimer or 3 µg B.1.351 S-Trimer (half adjuvanted and half non-adjuvanted). Animals in groups 2 to 3 were randomized to receive a booster (Dose 3) with either 3 µg of non-adjuvanted Prototype or B.1.351 S-Trimer. (**d**) BALB/c mice were immunized in Stage 1 with priming (Dose 1) on day 0 and boost (Dose 2) on day 21, with primary analysis for humoral immunogenicity was conducted on Day 35 blood samples. In Stage 2, a booster dose (Dose 3) was given on Day 35 and primary analysis for humoral and cellular immune responses was conducted on Day 49 blood samples.

Neutralizing antibody titers against B.1.351 and P.1 were highest in the groups receiving two doses of either B.1.351 S-Trimer and bivalent vaccine (Fig. 3A), with the former inducing the numerically-highest titers. Compared to animals receiving two doses of Prototype S-Trimer, animals receiving two doses of B.1.351 S-Trimer observed 13.8-fold higher neutralizing antibody titers against B.1.351 pseudovirus and 2.4-fold higher titers against P.1 pseudovirus. Interestingly, the group receiving heterologous prime boost (Dose 1 Prototype S-Trimer + Dose 2 B.1.351 S-Trimer) did not appear to induce higher antibody titers against B.1.351 and P.1 compared to animals receiving two doses of Prototype S-Trimer.

**Fig. 3.**
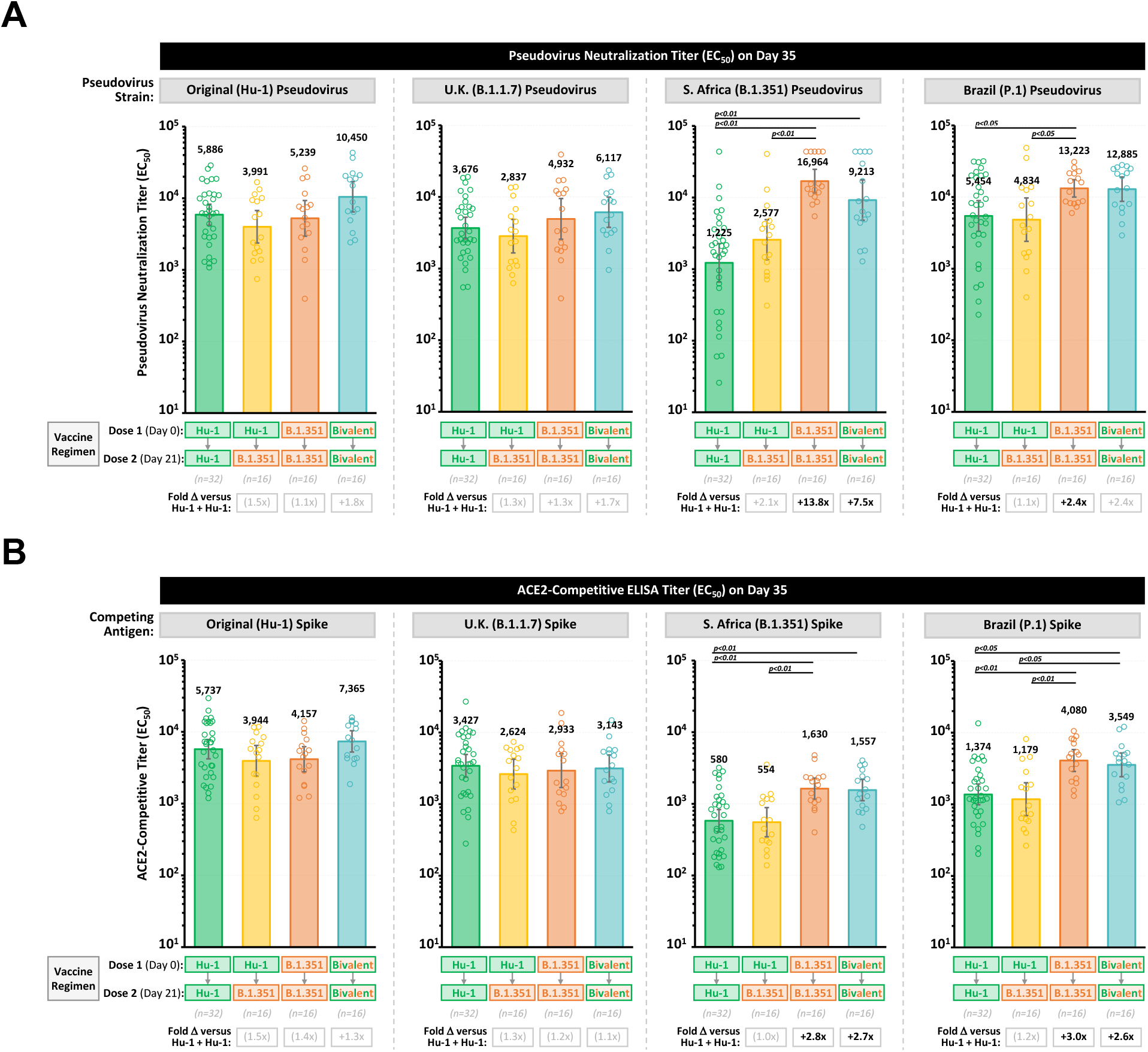
Humoral Immune Response of 2 Doses of Prototype and/or B.1.351 Spike-Trimer Antigens in Mice. Humoral immune responses in Stage 1 of the study were evaluated on Day 35 (2 weeks after Dose 2) based on (**a**) SARS-CoV-2 pseudovirus neutralization assays against original strain, UK (B.1.1.7), South African (B.1.351) and Brazil (P.1) strains and (**b**) ACE2-competitive ELISA detecting competition of vaccine-induced antibodies for binding to ACE2 with S-Trimers based on Original (Wuhan-Hu-1), UK (B.1.1.7), South African (B.1.351) and Brazil (P.1) strains. Dots represent individual animals; bars represent geometric mean titers (GMT) of half maximal effective concentration (EC_50_) values, and error bars represent 95% confidence intervals (95% CI). Fold differences (Δ) in GMTs for groups 2 to 4 compared to group 1 (Prototype S-Trimer) are shown, and statistically-significant differences are shown in black text. *P* values < 0.05 were considered significant.

Neutralizing antibody titers against the original and B.1.1.7 variant pseudoviruses were similar across all vaccine groups (Fig. 3A), demonstrating that two doses of the B.1.351 vaccine were able to elicit antibodies which are capable of fully back-neutralizing against the original strain and can also fully protect against the B.1.1.7 variant.

ACE2-competitive ELISA results were consistent with and further confirmed the pseudovirus neutralization results (Fig. 3B), and ACE2-competitive ELISA titers were found to be correlated with pseudovirus neutralization titers across all four strains tested (fig. S1).

The effect of adjuvanted versus non-adjuvanted second doses (boost) was done via factorial analysis based on pseudovirus neutralization titers (fig. S2A). Animals receiving an adjuvanted boost (antigen plus CpG 1018/alum) appeared to induce approximately 2- to 4-fold higher neutralizing antibody titers compared to animals receiving non-adjuvanted boost (antigen-only), demonstrating the positive impact of adjuvants on immune response when utilized for boost.

### Immunogenicity of Prototype or B.1.351 S-Trimer booster doses (dose 3) in mice

In Stage 2 of the study (Fig. 2C), animals in Prototype S-Trimer prime-boost group were randomized to receive a booster dose (Dose 3) on Day 35 with either Prototype S-Trimer or B.1.351 S-Trimer (half adjuvanted and half non-adjuvanted). Animals in the heterologous prime-boost group and B.1.351 S-Trimer prime-boost group were randomized to receive a booster dose (Dose 3) with either non-adjuvanted Prototype or non-adjuvanted B.1.351 S-Trimer. Analyses for humoral and cellular immune responses were conducted on Day 49 samples (2 weeks post-dose 3).

Across all vaccine groups after receiving a booster dose, neutralizing antibody titers against the original strain (Fig. 4A) and B.1.1.7 variant (Fig. 4B) were similar, and fold-increases in titers after the booster (Dose 3) compared to after Dose 2 were also similar across groups. Neutralizing antibody titers increased by about 1.6-to 3.1-fold against the original strain and 1.8- to 2.8-fold against B.1.1.7 variant on average across the booster groups.

**Fig. 4.**
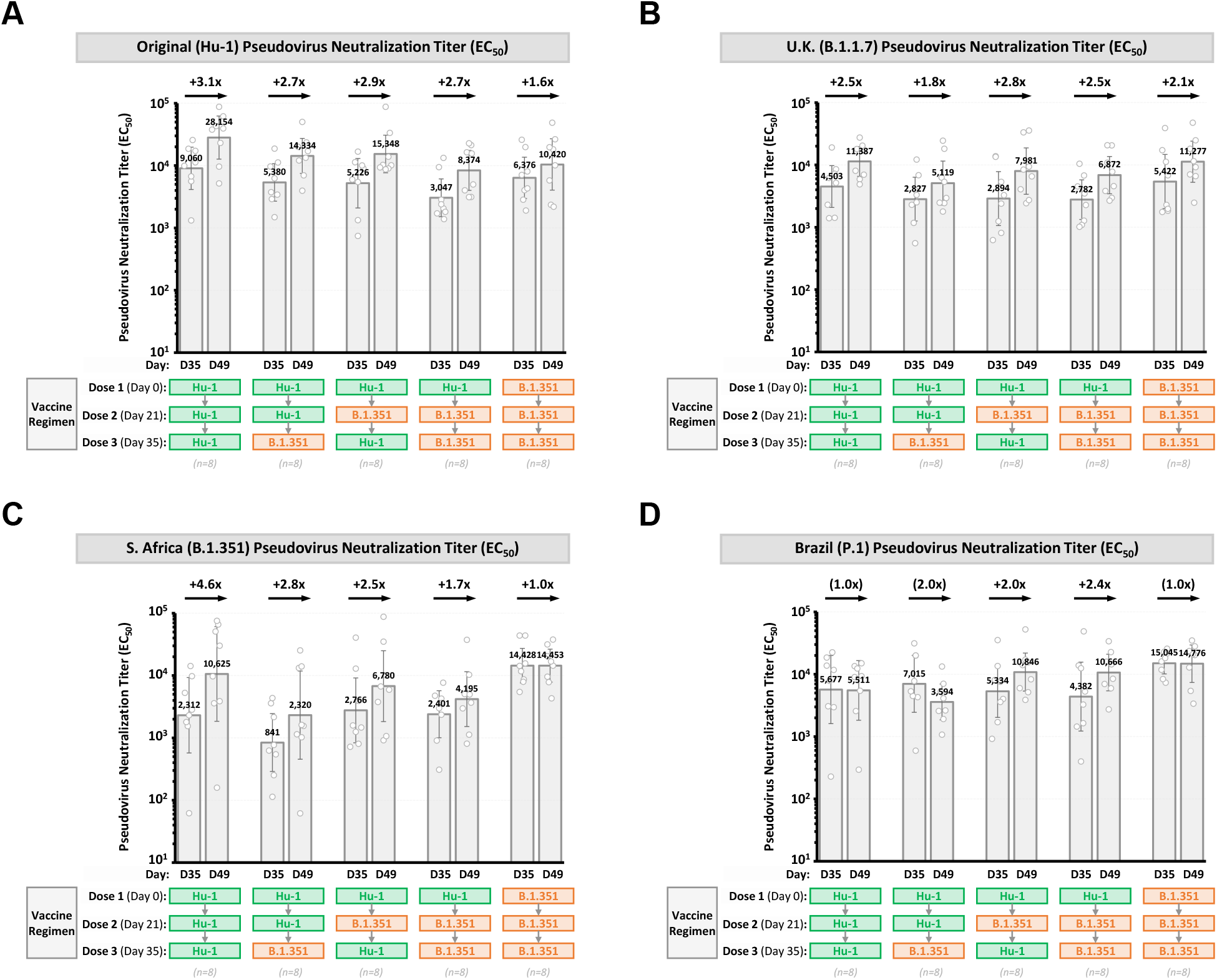
Humoral Immune Response of 3 Doses of Prototype and/or B.1.351 Spike-Trimer Antigens in Mice. Humoral immune responses in Stage 2 of the study were evaluated in animals receiving a non-adjuvanted booster (Dose 3) on Day 49 (2 weeks after Dose 3) based on SARS-CoV-2 pseudovirus neutralization assays against (**a**) Original (Wuhan-Hu-1) strain, (**b**) UK (B.1.1.7) variant, (**c**) South African (B.1.351) variant and (**d**) Brazil (P.1) variant pseudoviruses. Neutralization titers on Day 35 (2 weeks after Dose 2) are also shown. Fold differences in titers (Day 49 versus Day 35) in each booster group are shown. Dots represent individual animals; bars represent geometric mean titers (GMT) of half maximal effective concentration (EC_50_) values, and error bars represent 95% confidence intervals (95% CI).

In the groups receiving three doses of vaccine which included a Prototype S-Trimer priming dose (Dose 1), neutralizing antibody titers against the B.1.351 variant (Fig. 4C) increased by 1.7- to 4.6-fold on average after the booster dose, achieving levels comparable to neutralization against the original strain elicited by two doses of Prototype S-Trimer. However, we note that the variability in B.1.351 neutralization titers in these groups appeared to be higher than neutralization against the other strains tested. In the group receiving three doses of B.1.351 S-Trimer, neutralizing antibody titers against B.1.351 did not significantly increase after the booster dose, likely because titers were already at high biological levels after the boost (Dose 2).

Interestingly, in the groups receiving prime-boost with Prototype S-Trimer followed by a booster dose (either Prototype S-Trimer or B.1.351 S-Trimer), neutralization titers against the P.1 variant did not appear to increase following the booster dose (Fig. 4D). Similar to B.1.351 neutralization titers, P.1 neutralization titers were numerically highest in the group receiving three doses of B.1.351 S-Trimer.

We note that in the group of animals receiving two doses of Prototype S-Trimer vaccine (prime-boost), a subsequent booster dose (Dose 3) with B.1.351 S-Trimer did not appear to induce higher neutralizing antibodies against B.1.351 or P.1 compared to animals receiving a booster dose with Prototype S-Trimer (Fig. 4C, 4D).

The effect of adjuvanted versus non-adjuvanted booster doses (Dose 3) on humoral immunogenicity in group 1 was conducted using factorial analysis based on pseudovirus neutralization titers (fig. S2B). Animals receiving an adjuvanted booster dose (antigen plus CpG 1018/alum) did not appear to induce significantly higher neutralizing antibody titers compared to animals receiving non-adjuvanted booster doses (antigen-only).

### Cell-mediated immune response induced by 3 doses of Prototype or B.1.351 S-Trimer in mice

Cell-mediated immune responses were evaluated in animals receiving either three doses of Prototype S-Trimer or three doses of B.1.351 S-Trimer on Day 49 (2 weeks after dose 3). Cell-mediated immunity (CMI) was assessed by ELISpot detecting Th1 cytokines (IFNγ and IL-2) or Th2 cytokine (IL-5) in harvested mouse splenocytes stimulated with either original (Wuhan-Hu-1) strain S1 peptide pool, SARS-CoV S1 peptide pool, B.1.351 variant RBD peptide pool, or P.1 variant RBD peptide pool.

Strong Th1-biased CMI was observed in both vaccine groups across all stimulants tested (Fig. 5A-D). The Th1-biased CMI induced by adjuvanted S-Trimer antigen is consistent with results from our previous studies (*16*).

**Fig. 5.**
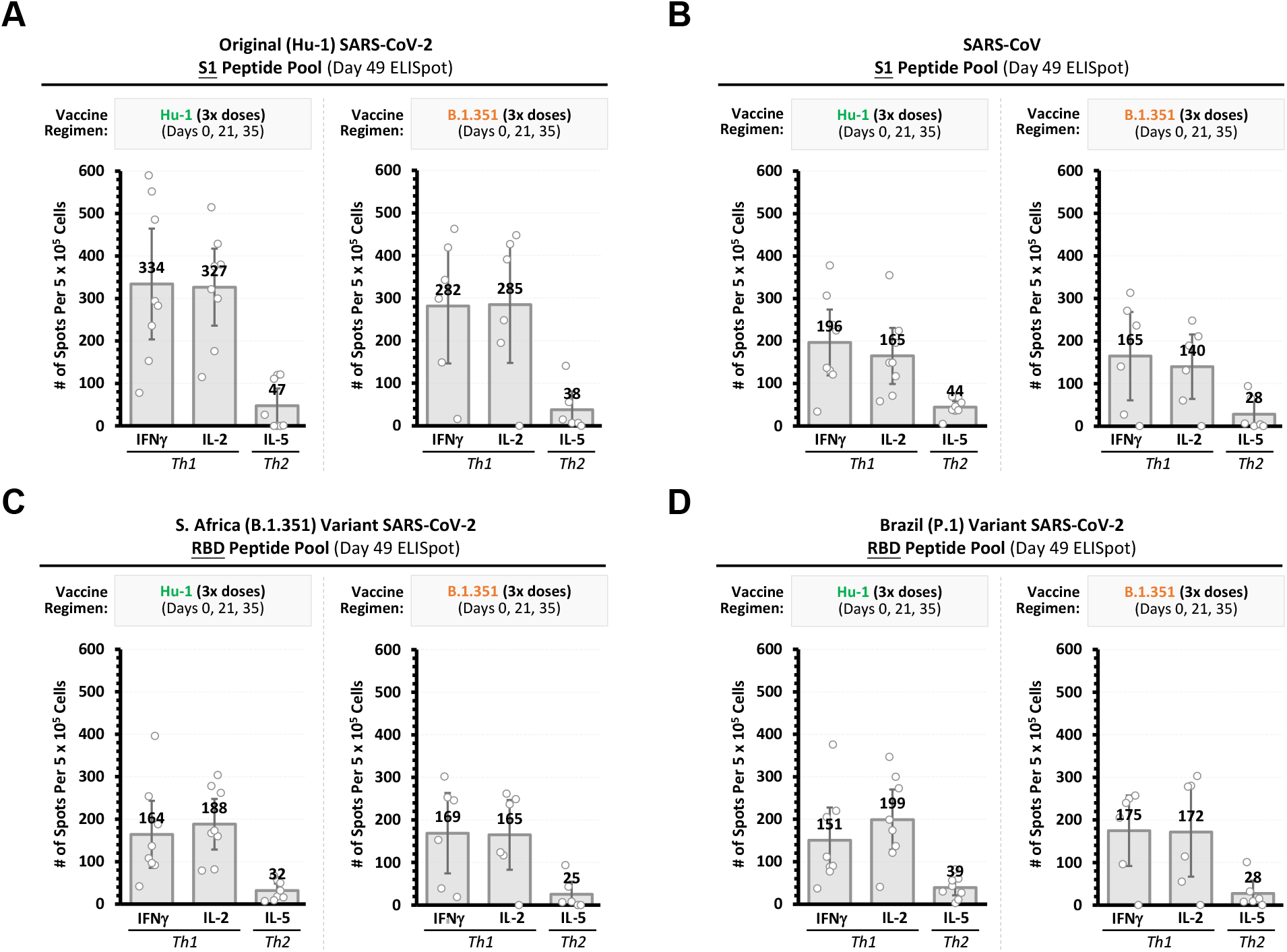
Cell-Mediated Immune Responses of 3 Doses of Prototype or B.1.351 Spike-Trimer Antigens in Mice. Cell-mediated immune responses in Stage 2 of the study were evaluated on Day 49 (2 weeks after dose 3) based on ELISpot detecting Th1 cytokines (IFNγ and IL-2) or Th2 cytokine (IL-5) in harvested splenocytes stimulated with (**a**) S1 peptide pool from original (Wuhan-Hu-1) strain, (**b**) S1 peptide pool from SARS-CoV, (**c**) RBD peptide pool from B.1.351 variant, or (**d**) RBD peptide pool from P.1 variant. Dots represent individual animals; bars represent group mean values, and error bars represent 95% confidence intervals (95% CI).

Importantly, the magnitude of CMI appeared similar in both vaccine groups across all stimulants, suggesting that broad cross-reactive CMI directed to VOCs (including B.1.1.7, B.1.351 and P.1) can be induced by both the Prototype and B.1.351 S-Trimer vaccines. Cross-reactive CMI against SARS-CoV S1 peptide pool (Fig. 5B) did appear to be at levels which were about 40 to 50% lower than CMI to original strain of SARS-CoV-2 (Fig. 5A), in-line with the 79% sequence homology of the S1 domains for SARS-CoV and SARS-CoV-2.

## Discussion

### Modified B.1.351 S-Trimer vaccine induces broad neutralization against original strain and variants of concern

Consistent with results from other studies evaluating prototype COVID-19 vaccines based on the original strain of SARS-CoV-2 (*5,8,9,10*), we observed in this study that Prototype S-Trimer prime-boost vaccination induced lower neutralizing antibody titers against B.1.351 and P.1 variants compared to titers against the homologous original SARS-CoV-2 strain (Fig. 3). It is believed that the E484K and K417 mutations in the RBD of both B.1.351 and P.1 variants confer immune escape from antibodies induced by prototype vaccines (*3*).

As expected, two doses of our modified B.1.351 S-Trimer antigen induced high levels of neutralizing antibodies against the B.1.351 and P.1 variants. Importantly, two doses of B.1.351 S-Trimer was also able to fully back-neutralize against the original SARS-CoV-2 strain, as well as neutralize the B.1.1.7 variant. In contrast, an mRNA COVID-19 vaccine candidate based on the B.1.351 variant spike protein (containing all mutations found in B.1.351) appeared to induce approximately 6-fold lower neutralizing antibody titers against the original SARS-CoV-2 strain compared to the prototype mRNA vaccine in mice (*21*).

The broad neutralization induced by our modified B.1.351 S-Trimer vaccine candidate could potentially be explained by the chimeric nature of the antigen sequence, which contains 3 RBD mutations (K417N, E484K, N501Y) and the D614G mutations found in the B.1.351 variant strain, while the NTD and S2 sequences are based on the original SARS-CoV-2 strain (Wuhan-Hu-1) (Fig. 1a). Previous studies have identified neutralizing antibodies that target NTD (*22,23*), and emerging evidence suggests that vaccine-induced anti-NTD neutralizing antibodies are co-dominant with anti-RBD neutralizing antibodies (*24*), highlighting the importance of NTD. Thus, the data suggests a potential model for inducing broad neutralization whereby our modified B.1.351 S-Trimer antigen could induce (a) anti-RBD antibodies targeting E484K and K417 mutations that can neutralize B.1.351 and P.1 variant strains and (b) anti-NTD antibodies that can neutralize the original strain.

However, a study in COVID-19 convalescent plasma from patients infected with E484K-containing virus strains appeared to observe neutralizing antibodies against the original strain at levels comparable to patients who were infected with the original strain virus (*25*).

Thus, additional studies are needed to confirm the nature of the broad neutralization induced by our modified B.1.351 S-Trimer vaccine candidate. Studies in non-human primates to confirm the broadly neutralizing characteristics of the modified B.1.351 vaccine are planned, as well as studies comparing the immunogenicity of the modified B.1.351 S-Trimer vaccine with an S-Trimer containing all mutations found in the B.1.351 variant strain spike protein (including NTD and S2).

### Booster doses following two doses of prototype vaccine could strengthen broad neutralization

Given the global rollout of prototype COVID-19 vaccines (based on the original SARS-CoV-2 strain) starting in early 2021 (*13*) and the apparent lower levels of neutralization and vaccine efficacy induced by these vaccines against VOCs, an important question is if the administration of additional booster doses can increase cross-neutralization against VOCs to highly-protective levels. If so, then should booster vaccine doses be based on variants or can prototype vaccines also be used?

Results in this study suggest that a booster dose (with either Prototype S-Trimer or B.1.351 S-Trimer) following two doses of Prototype S-Trimer could increase levels of broad neutralizing antibodies against the original strain and VOCs, and neutralizing antibody titers against B.1.351 increased to levels comparable to neutralizing antibody titers against the original strain elicited by two doses of Prototype S-Trimer. There did not appear to be significant differences in neutralizing antibody titers if the booster dose given was Prototype S-Trimer or B.1.351 S-Trimer. These findings appear to be consistent with preliminary Phase 1 clinical trial data evaluating booster doses of mRNA vaccine candidates based on either the original strain or B.1.351 variant (*26*). If these findings are further evaluated and confirmed in humans, greater flexibility in effective boosting strategies could be employed based on the supply of prototype and variant vaccines available at a given time. However, we note that the longer-term effects of continued boosting or re-vaccination with either homologous or heterologous antigens remain to be studied.

It should also be noted that heterologous prime-boost (Dose 1 Prototype S-Trimer + Dose 2 B.1.351 S-Trimer) did not induce higher neutralizing antibody titers against B.1.351 and P.1 compared to animals receiving homologous prime-boost with Prototype S-Trimer. Furthermore, while a booster dose following two doses of Prototype S-Trimer increased B.1.351 neutralization titers by about 2.8- to 4.6-fold (compared to post-dose 2 levels), the titers appeared to be more variable and numerically lower than animals receiving two doses or three doses of B.1.351 S-Trimer. Additionally, P.1 neutralization titers did not increase in these groups which received Prototype S-Trimer prime-boost followed by a booster and remained 2.7- to 4.1-fold lower than the group receiving three doses of B.1.351 S-Trimer. These results suggest that priming with prototype COVID-19 vaccines may induce some degree of ‘original antigenic sin’, a phenomenon previously-described for other viruses (such as influenza and dengue) whereby the immune system preferentially responds to the epitopes in the priming immunogen and is less able to respond to the new epitopes in the variant (*27*). However, as noted above, cross-reactive neutralizing antibody titers against VOCs in mice induced by Prototype S-Trimer prime-boost were still relatively high, albeit lower than levels induced by the B.1.351 S-Trimer, and neutralization titers could be increased with a booster dose.

The encouraging results of S-Trimer booster doses in this study also warrant the further evaluation of heterologous prime-boost strategies across different platforms, utilizing our protein-based S-Trimers a booster dose following primary vaccination series with adenovirus-vectored, mRNA, or inactivated COVID-19 vaccines. Some concerns have emerged around the safety of multiple doses of mRNA vaccines including potential polyethylene glycol (PEG) sensitization (*28*), and anti-vector immunogenicity for adenovirus-vectored vaccines are known to reduce the effectiveness of subsequent homologous doses (*29,30,31*). Preliminary human clinical data indicates that heterologous prime-boost with adenovirus-vectored and mRNA COVID-19 vaccines (or vice versa) suggest that they induce higher rates of systemic reactogenicity than heir homologous prime-boost counterparts (*32*), further warranting the evaluation of protein-based candidates as booster doses.

### Broad cross-reactive cell-mediated immune responses against SARS-CoV-2

While the precise role of cell-mediated immunity (CMI) in the prevention or recovery from COVID-19 remains to be elucidated, there is ample evidence that strong cellular immune responses against the spike (S) protein are induced in COVID-19 patients (*33*) and that CMI could contribute to the attenuation of symptoms and accelerated clearance of SARS-CoV-2 (*34,35*). Furthermore, it has been reported that up to ∼35% of SARS-CoV-2 naïve individuals have some degree of cross-reactive CD4^+^ T-cell responses to SARS-CoV-2 antigens due to prior infection by other common-cold coronaviruses (*36*). In this study, while lower levels of neutralizing antibodies against B.1.351 were induced by Prototype S-Trimer compared with B.1.351 S-Trimer, CMI levels appeared to be similar for both vaccine candidates and were cross-reactive for antigens to SARS-CoV-2 (original strain S1, B.1.351 variant RBD, P.1 variant RBD) and SARS-CoV (S1), suggesting that CMI against coronavirus spike proteins could be more broadly cross-reactive than humoral immune responses.

Encouragingly, the CMI induced by the S-Trimers in this study appeared to be Th1-biased in nature, which is consistent with our previous studies evaluating adjuvanted Prototype S-Trimer (*16,17*).

### Use of adjuvants in boosting strategies

Given the high productivity of S-Trimer antigens utilizing Trimer-Tag technology and the potential to rapidly scale-up production to billions of doses (*16*), we evaluated the potential for ‘adjuvant sparing’ in this study should the supply of adjuvants be a limiting factor for the availability of our COVID-19 vaccines.

Because the use of adjuvants is believed to be most important for their role in establishing immunological priming (*37,38*), we evaluated and compared the administration of adjuvanted versus non-adjuvanted second doses (boost) and third doses (booster) doses. Animals receiving an adjuvanted boost (antigen plus CpG 1018/alum) induced 2- to 4-fold higher neutralizing antibody titers compared to animals receiving non-adjuvanted boost (antigen-only) (fig. S2A), suggesting that adjuvants may still be needed for second doses in the primary vaccination series to achieve an optimal immune response. However, animals receiving an adjuvanted booster (third dose) did not appear to induce significantly higher neutralizing antibody titers compared to animals receiving non-adjuvanted booster doses (antigen-only) (fig. S2B). These results warrant the further evaluation of both adjuvanted and non-adjuvanted booster strategies, and evaluation of boost/booster doses adjuvanted with alum-alone is also planned.

## Methods

### Animal studies, facilities and ethics statements

Specific pathogen-free (SPF) BALB/c female mice (6-8 weeks old) for immunogenicity studies were purchased from Beijing Vital River Laboratory Animal Technology Co., Ltd. and kept under standard pathogen-free conditions in the animal care center at Chengdu Hi-tech Incubation Park. All animals were allowed free access to water and diet and provided with a 12 h light/dark cycle (temperature: 16-26°C, humidity: 40%-70%). All mouse experiments were approved by the institutional animal care and use committee (IACUC) in Clover Biopharmaceuticals and were conducted according to international guidelines for animal studies.

### Human COVID-19 convalescent serum samples

Human convalescent sera samples from recovered COVID-19 patients (table S1) infected with the original SARS-CoV-2 strain were obtained from Public Health Clinical Center of Chengdu in Chengdu, China, under approved guidelines by the Institutional Review Board (IRB), and all patients had provided written informed consent before sera sample were collected. All convalescent sera samples were heat inactivated at 55°C for 30 min before being used for analysis.

### Adjuvants

CpG 1018 (*39*) was manufactured under GMP by Dynavax Technologies. CpG 1018, a TLR-9 agonist, is a synthetic CpG-B class oligonucleotide having a phosphorothioate-backbone and the sequence 5′-TGACTGTGAA<u>CG</u>TT**CG**AGATGA-3′. Alum hydroxide was manufactured under GMP by Croda. The S-Trimer subunit vaccines were mixed with the adjuvants by gentle inversion in 1:1 ratio by volume preceding each immunization.

### Protein expression and purification

The Prototype S-Trimer and modified B.1.351 S-Trimer antigens were produced and purified as previously described (*16*) and used for immunogenicity studies. S-Trimer antigens based on B.1.1.7 variant and P.1 variant were also produced and used for ACE2-competitive ELISA assays. To rapidly express the variant S-Trimer antigens which are covalently-trimerized, we employed Trimer-Tag technology (*16*). cDNA encoding the ectodomain of the respective SARS-CoV-2 variant strain Spike (S) proteins were subcloned into the pTRIMER mammalian expression vector to allow in-frame fusion to Trimer-Tag, which is capable of self-trimerization via disulfide bonds. After transient transfection in 293F cells, the variant S-Trimer antigens were expressed and secreted at sufficient levels to enable further characterization and mouse immunogenicity studies. To obtain the variant S-Trimer antigens in a highly-purified form for characterization and vaccine studies, we utilized an affinity purification scheme as previously described (*16*), allowing us to purify the antigens to near homogeneity in a single step.

### Receptor binding studies of S-Trimer to human ACE2

The binding affinity of S-Trimer to ACE2 was assessed by Bio-Layer Interferometry measurements on ForteBio Octet QKe (Pall). ACE2-Fc (10 µg/mL) was immobilized on Protein A (ProA) biosensors (Pall). Real-time receptor-binding curves were obtained by applying the sensor in two-fold serial dilutions of S-Trimer (2.25-36 µg/mL in PBS). Kinetic parameters (K_on_ and K_off_) and affinities (K_D_) were analyzed using Octet software, version 12.0. Dissociation constants (K_D_) were determined using steady state analysis, assuming a 1:1 binding model for a S-Trimer (Prototype or B.1.351) to ACE2-Fc.

### Immunogenicity analysis of Prototype S-Trimer and modified B.**1.351 S-Trimer in mice**

BALB/c mice (n=16-32/group) were immunized in Stage 1 of the study (Fig. 2C) with either two doses (Day 0 and Day 21) of Prototype S-Trimer (3 µg), heterologous prime-boost (dose 1 Prototype S-Trimer; dose 2 B.1.351 S-Trimer; 3 µg of each antigen), two doses B.1.351 S-Trimer (3 µg), or two doses of bivalent vaccine (3 µg Prototype S-Trimer mixed with 3 µg B.1.351 S-Trimer). All animals in Stage 1 received a priming dose (Dose 1) adjuvanted with 10 µg CpG 1018 (Dynavax) plus 75 µg Alum (aluminum hydroxide, Croda), whereas half of the animals in each group received a boost (Dose 2) adjuvanted with CpG 1018 plus Alum, and the other half received non-adjuvanted boost (antigen-only). In Stage 2 (Fig. 2C) of the study, animals in group 1 were randomized to receive a booster (Dose 3 on Day 35) with either 3 µg Prototype or 3 µg B.1.351 S -Trimer (half adjuvanted and half non-adjuvanted). Animals in groups 2 to 3 were randomized to receive a booster (Dose 3) with either 3 µg of non-adjuvanted Prototype or B.1.351 S-Trimer. In Stage 1, primary analysis for humoral immunogenicity was conducted on Day 35 blood samples. In Stage 2, primary analysis for humoral and cellular immune responses was conducted on Day 49 blood samples. Animals were bled from tail veins for humoral immune responses analyses. Spleens were removed after sacrifice for ELISpot assays.

### ACE2-competitive ELISA assays

96-well plates (Corning) were coated with 1 μg/mL ACE2-Fc (100 μL/well) at 4°C overnight, blocked with 2% non-fat milk 37°C for 2 h. After washing 3 times with PBST, the plates were incubated with 100 ng/mL S-Trimer (Prototype, B.1.351, B.1.1.7 or P.1) mixed with serially diluted antisera for 1 h at 37°C. After washing 3 times with PBST, the plates were incubated with 1:5000 dilution of rabbit anti-Trimer-Tag antibody (Clover Biopharmaceuticals) at 37°C for 1 h, followed by washing 3 times with PBST and then a 1:20000 dilution of goat anti-rabbit IgG-HRP (Southern Biotech). After washing 3 times with PBST, TMB (Thermo Scientific) was added for signal development. The percentage of inhibition was calculated as follows: % inhibition = [(A-Blank)-(P-Blank)]/(A-Blank)x100, where A is the maximum OD signal of S-Trimer binding to ACE2-Fc when no serum was present, and P is the OD signal of S-Trimer binding to ACE2-Fc in presence of serum at a given dilution. The EC_50_ of a given serum sample was defined as the reciprocal of the dilution where the sample shows 50% competition.

### Pseudovirus neutralization assays

SARS-CoV-2 pseudovirus neutralization assay for the original (Wuhan-Hu-1) strain and variant B.1.1.7, B.1.351 and P.1. strains (Beijing Tiantan Pharmaceutical Biotechnology Development Co. Ltd.) were conducted as previously described (*40*), with some modifications. To evaluate the SARS-CoV-2 pseudovirus neutralization activity of antisera, samples were first heat-inactivated for 30 min and serially diluted (3-fold), incubated with an equal volume of 650 TCID_50_ pseudovirus at 37°C for 1 h, along with virus-alone (positive control) and cell-alone (negative control). Then, freshly-trypsinized ACE2 overexpression-293 cells were added to each well at 20000 cells/well. Following 24 h incubation at 37°C in a 5% CO2 incubator, the cells were lysed and luciferase activity was determined by a Luciferase Assay System (Beyotime), according to the manufacturer’s protocol. The EC_50_ neutralizing antibody titer of a given serum sample was defined as the reciprocal of the dilution where the sample showed the relative light units (RLUs) were reduced by 50% compared to virus alone control wells.

### Splenocyte stimulation and ELISpot assays

To detect antigen-specific T-cell responses, ELISpot kits (Mabtech) measuring Th1 cytokines (IFN-γ, IL-2) and Th2 cytokine (IL-5) were used per manufacturer’s instructions. Splenocytes from immunized mice or PBMC from immunized mice were harvested 2 weeks after the third immunization. 5 × 10^5^ splenocytes (96-well plate) were stimulated *in vitro* with 2 µg/mL of either original SARS-CoV-2 S1 peptide pool, SARS-CoV S1 peptide pool, B.1.351 RBD peptide pool or P.1 RBD peptide pool. Phorbol 12-myristate 13-acetate (PMA) and ionomycin as the non-specific stimulus were added to the positive control wells, whereas the negative control well received no stimuli. After 24-48 h incubation, biotinylated detection antibodies from the ELISpot kits and SA-ALP/SA-HRP were added. Blots were developed by the addition of BCIP/NBT or AEC substrate solution, which produced colored spots after 5-30 min incubation in the dark. Finally, the IFN-γ, IL-2 and IL-5 spot-forming cells (SFCs) were counted using an automatic ELISpot reader (CTL).

### Statistical analysis

Data arrangement was performed by Excel and statistical analyses were performed using the Prism 8.0 (GraphPad Software). Comparisons among multiple groups were performed using Kruskal-Wallis ANOVA with Dunn’s multiple comparisons tests. Factorial analysis was employed for certain analyses. P values < 0.05 were considered significant. ns, no significance.

## ACKNOWLEDGMENTS

The authors would like to acknowledge and thank the Clover Scientific Advisory Board (Donna Ambrosino, Sue Ann Costa Clemens, Pierre Desmons, Sam Liao, Michael Pfleiderer, Antoinette Quinsaat, Frank Rockhold, David Salisbury, George Siber, Nelson Teich, Anh Wartel, and Nicholas Jackson) for helpful expert advice and support as well as critical review and comments for this manuscript. The authors would like to thank Dynavax Technologies Corporation for providing CpG 1018 adjuvant for this study.

## Author Contribution

J.G.L and P.L. conceived this project, and J.G.L and D.S. designed the study. D.S. oversaw mouse studies, cell culture for antigen production and developed *in vitro* antibody/neutralizing antibody assays. X.L and C.H. performed expression vector construction and antibody titer experiments. P.Luo and X.Y. conducted protein purification experiments. R.X. and J. Wu directed project management for parts of this study. Y.L directed quality control experiments. Y.L. and X.H. performed binding affinity experiments. Q.W. and W.Q. performed the animal studies. X.H and M.C. performed cell-mediated immune response experiments. Y.Z., M.L. and D.L. collected and provided human convalescent sera for this study. J.G.L., D.S., R.C., G.S., D.A., D.M.S. analyzed the data and provided critical comments for this manuscript. J.G.L. and D.S. wrote the manuscript with input from all other authors.

## Supplementary Figures

**Fig. S1.**
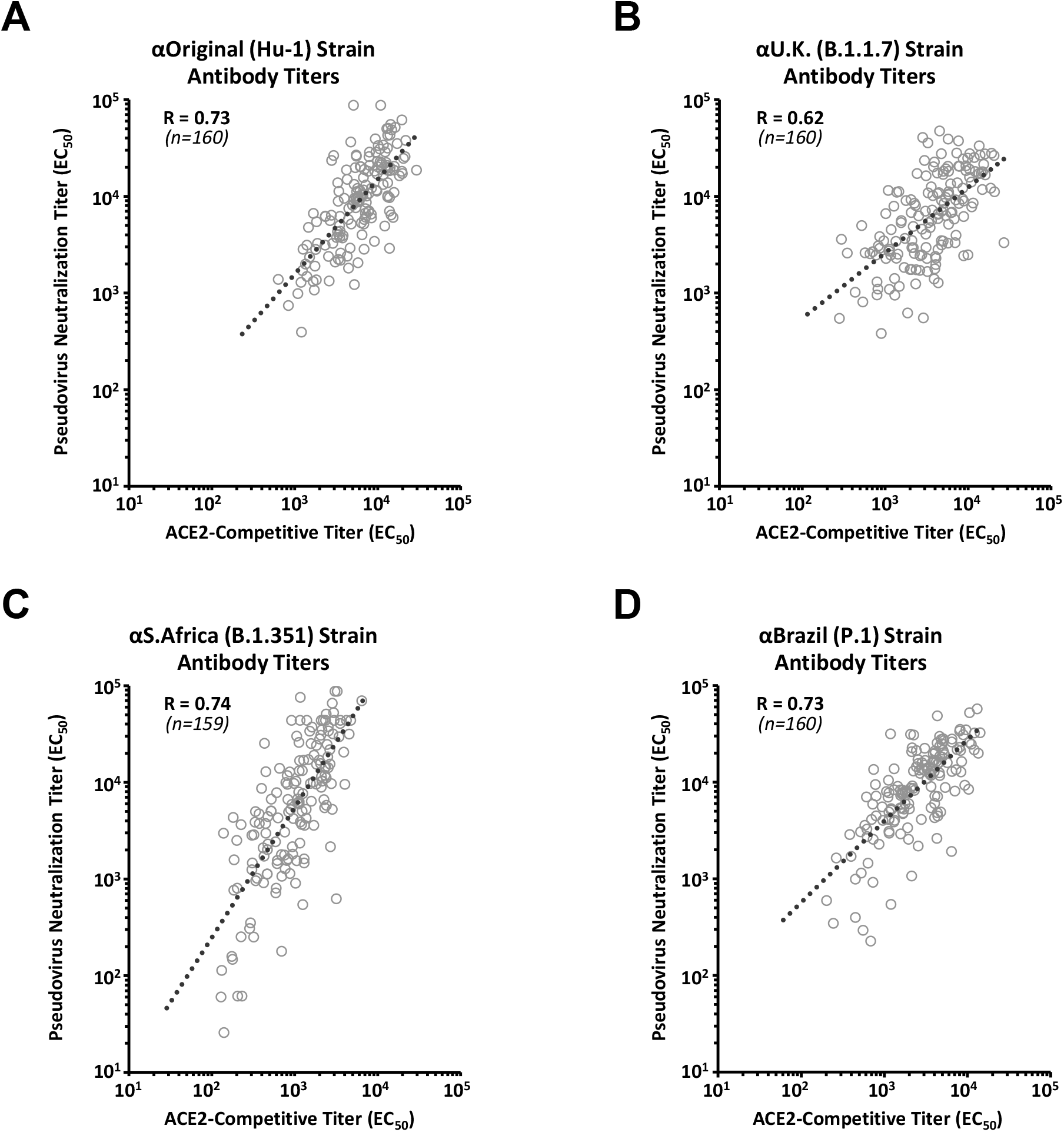
Correlation of antibody titers in immunized mice. Antibody titers in immunized mice based on pseudovirus neutralization assay and ACE2-competitive ELISA specific for (**a**) Original strain, (**b**) UK (B.1.1.7) variant, (**c**) South African (B.1.351) variant and (**d**) Brazil (P.1) variant strains were analyzed for correlation based on two-tailed Pearson’s R analysis. Data presented are pooled from post-immunization samples at Day 35 and Day 49. Points represent individual animal samples.

**Fig. S2.**
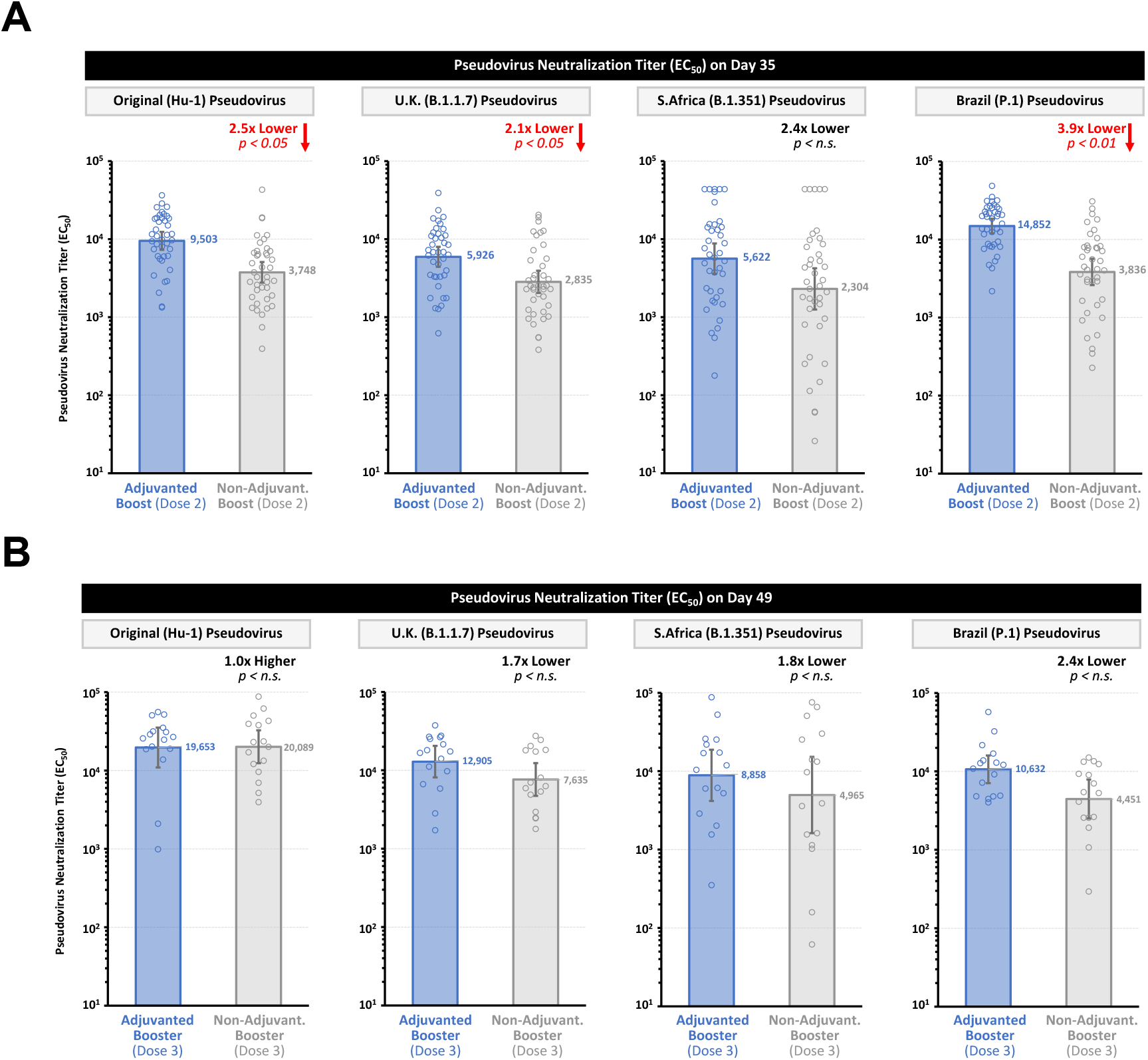
Effect of Adjuvants on Humoral Immune Response for Boost (Dose 2) and Booster (Dose 3). Humoral immune responses in the study were evaluated based on SARS-CoV-2 pseudovirus neutralization assays against original strain, UK (B.1.1.7), South African (B.1.351) and Brazil (P.1) strains. Results for pseudovirus neutralization titers based on factorial analyses and are shown here for (**a**) Stage 1 where all animals received a priming dose (Dose 1) adjuvanted with CpG 1018 plus alum, whereas half of the animals received an adjuvanted boost (Dose 2) with CpG 1018 plus alum, and the remaining half of the animals received non-adjuvanted boost (antigen-only), and (**b**) Stage 2 where half of the animals in Group 1 received an adjuvanted booster Dose 3 (CpG 1018 plus alum) and the remaining half of the animals received non-adjuvanted booster (antigen-only). Dots represent individual animals; bars represent geometric mean titers (GMT) of EC_50_ values, and error bars represent 95% confidence intervals (95% CI). *P* values < 0.05 were considered significant.

**Fig. S3.**
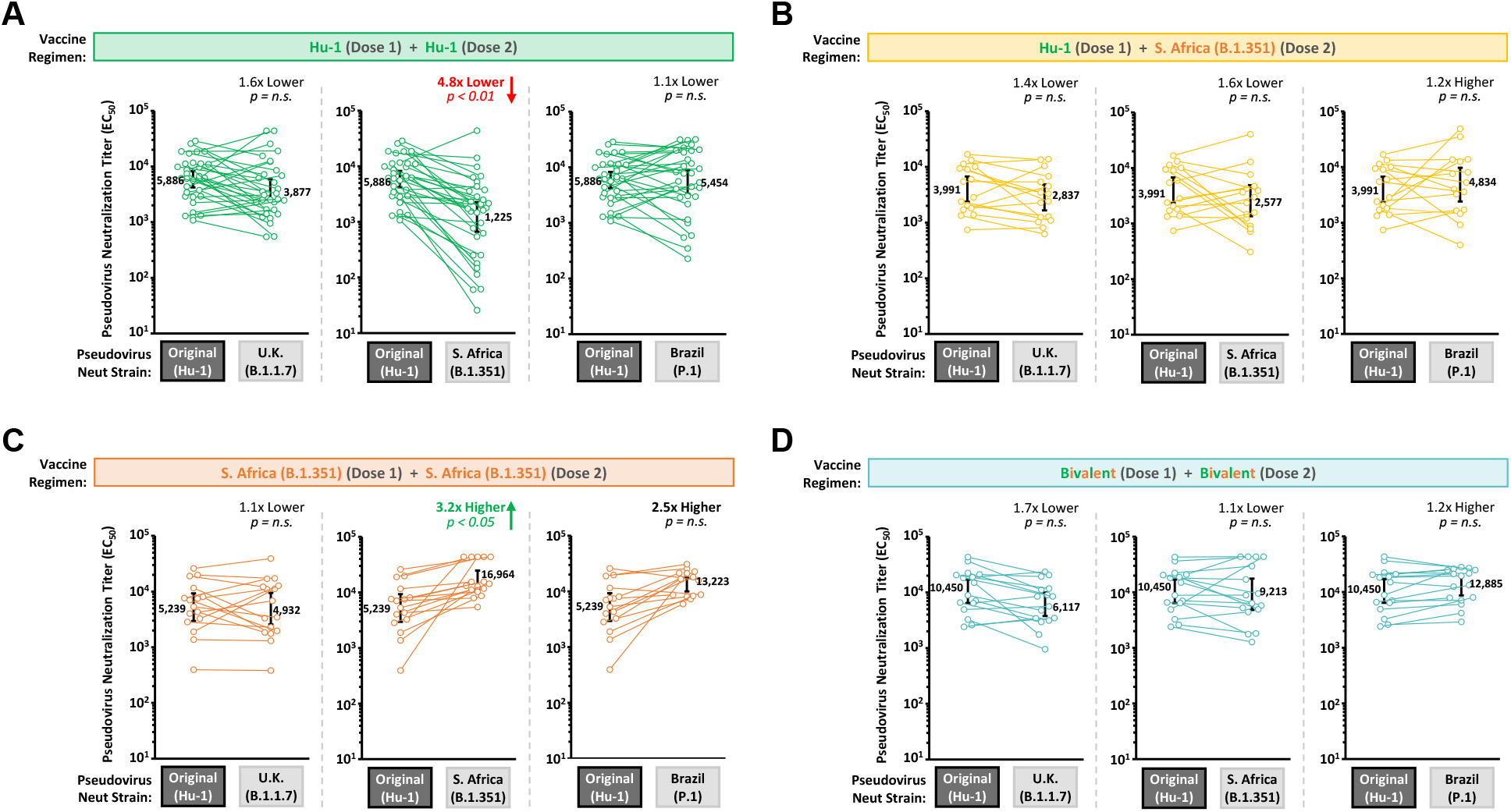
Cross-Neutralization Based on Pseudovirus Neutralization Titers After 2 Doses of Prototype and/or B.1.351 S-Timer Antigens in Mice. Humoral immune responses in Stage 1 of the study were evaluated on Day 35 (2 weeks after dose 2) based on SARS-CoV-2 pseudovirus neutralization assays against original strain, UK variant (B.1.1.7), South African variant (B.1.351) and Brazil variant (P.1) strains. Results are shown for (**a**) Group 1 (two doses of Prototype S-Trimer), (**b**) Group 2 (heterologous prime-boost), (**c**) Group 3 (two doses B.1.351 S-Trimer), or (**d**) two doses of bivalent vaccine. Results from individual animals are represented by dots in each figure, with lines connecting the Original and variant neutralization titers. Geometric mean titers (GMT) of EC_50_ values are shown, and error bars represent 95% confidence intervals (95% CI). *P* values < 0.05 were considered significant (n.s., not significant).

**Fig. S4.**
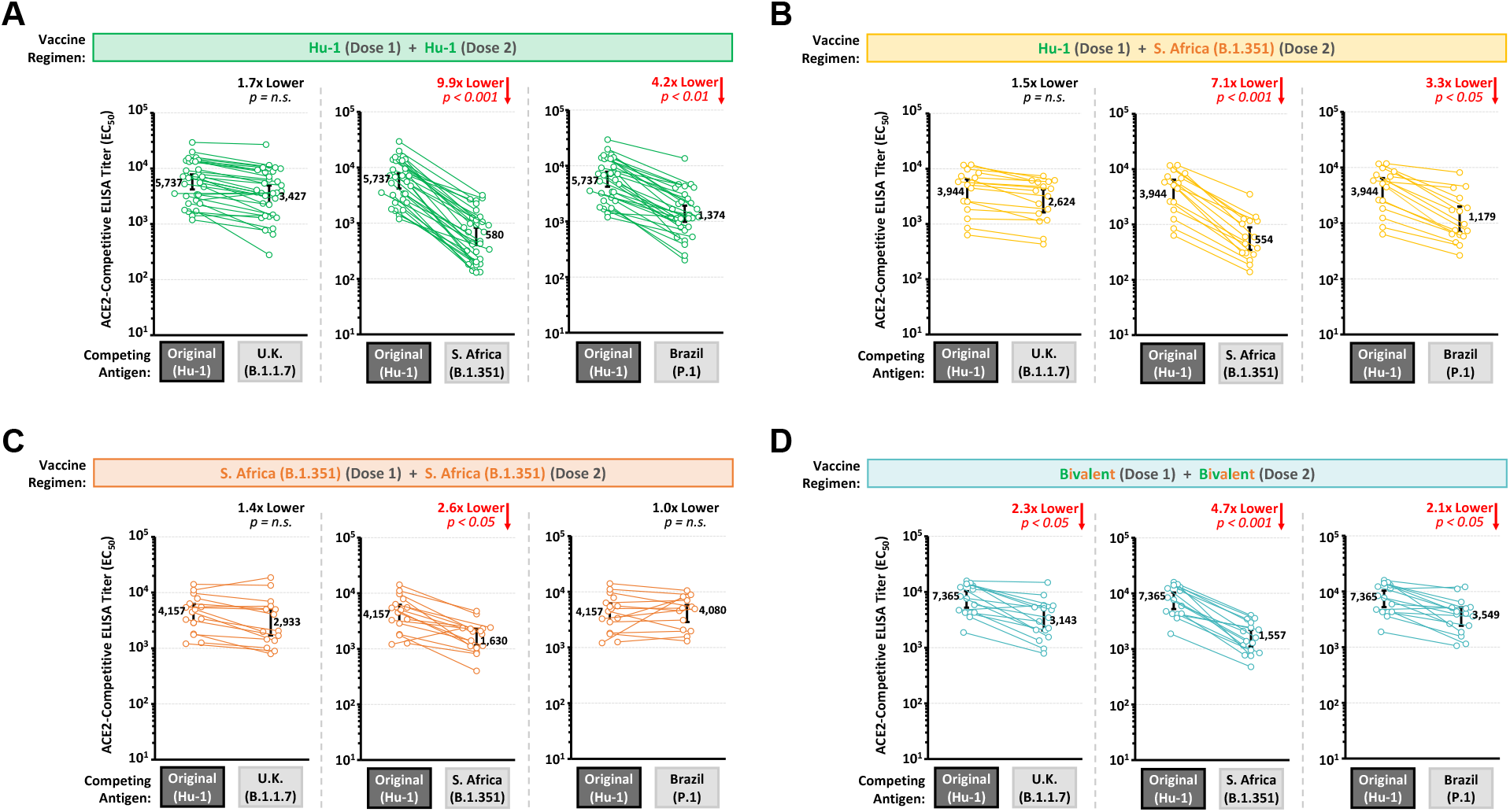
Cross-Neutralization Based on ACE2-Competitive Titers After 2 Doses of Prototype and/or B.1.351 S-Timer Antigens in Mice. Humoral immune responses in Stage 1 of the study were evaluated on Day 35 (2 weeks after dose 2) based on ACE2-competitive ELISA assays against original strain, UK variant (B.1.1.7), South African variant (B.1.351) and Brazil variant (P.1) strains. Results are shown for (**a**) Group 1 (two doses of Prototype S-Trimer), (**b**) Group 2 (heterologous prime-boost), (**c**) Group 3 (two doses B.1.351 S-Trimer), or (**d**) two doses of bivalent vaccine. Results from individual animals are represented by dots in each figure, with lines connecting the Original and variant neutralization titers. Geometric mean titers (GMT) of EC_50_ values are shown, and error bars represent 95% confidence intervals (95% CI). *P* values < 0.05 were considered significant (n.s., not significant).

**Fig. S5.**
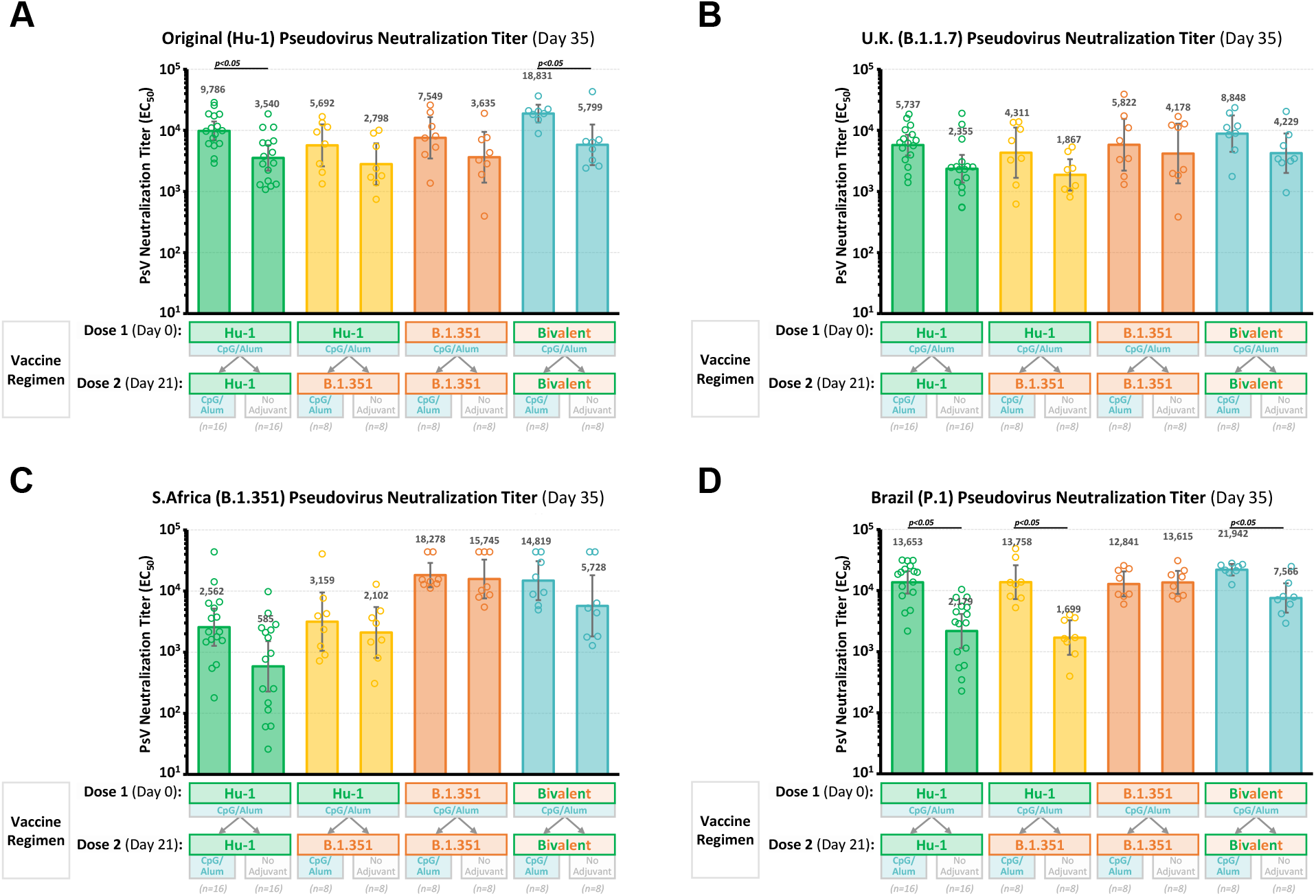
Humoral Immune Response After 2 Doses of Prototype and/or B.1.351 S-Timer Antigens in Mice. Humoral immune responses in Stage 1 of the study were evaluated on Day 35 (2 weeks after dose 2) based on SARS-CoV-2 pseudovirus neutralization assays against (**a**) original strain, (**b**) UK variant (B.1.1.7), (**c**) South African variant (B.1.351) and (**d**) Brazil variant (P.1) strains. All animals in Stage 1 received a priming dose (Dose 1) adjuvanted with CpG 1018 plus alum, whereas half of the animals in each group received a boost (Dose 2) adjuvanted with CpG 1018 plus Alum, and the other half received non-adjuvanted boost (antigen-only). Results here are shown for subgroups of animals receiving either adjuvanted or non-adjuvanted boost (dose 2). Dots represent individual animals; bars represent geometric mean titers (GMT) of EC_50_ values, and error bars represent 95% confidence intervals (95% CI). *P* values < 0.05 were considered significant.

**Fig. S6.**
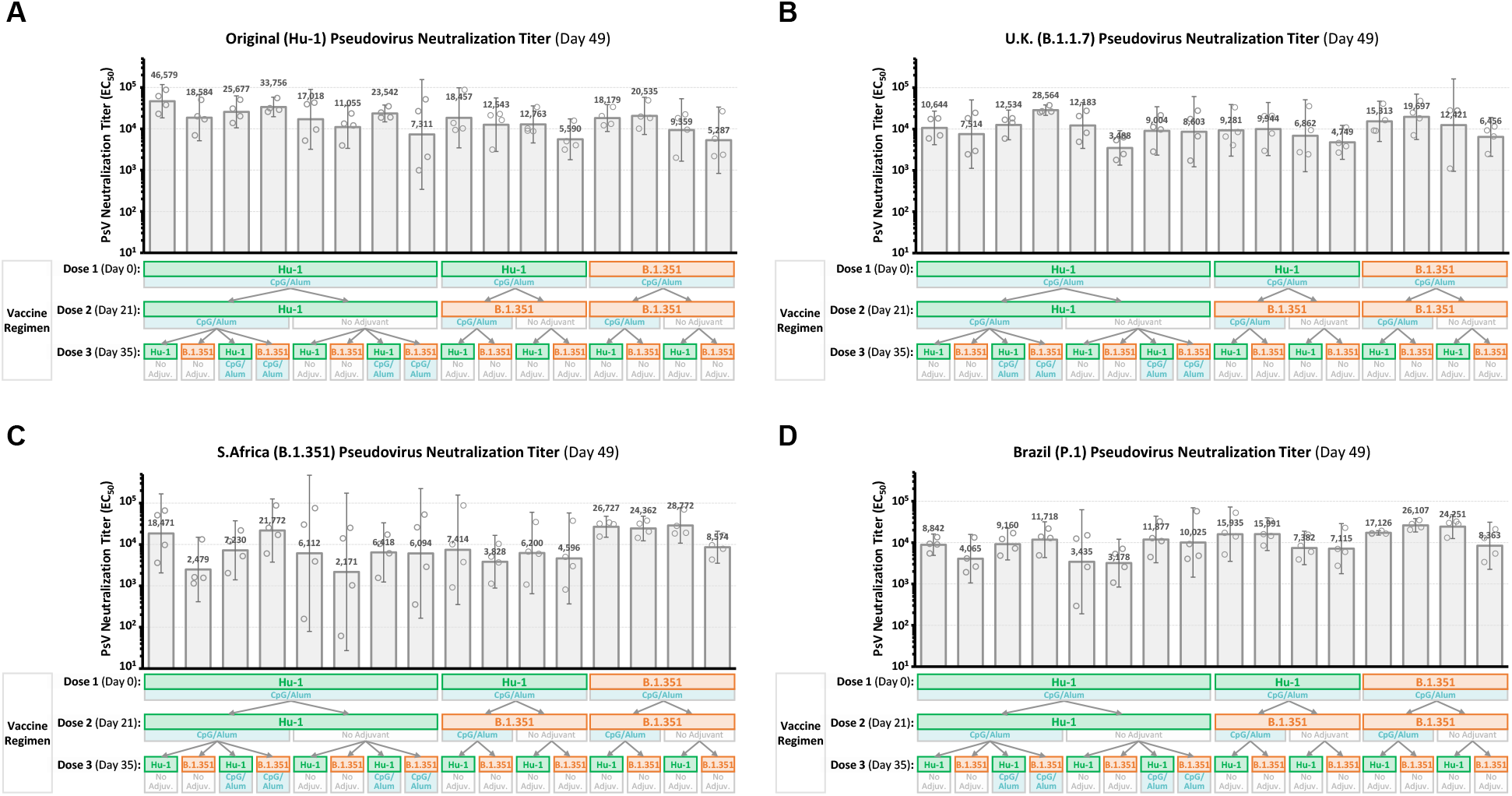
Humoral Immune Response After 3 Doses of Prototype and/or B.1.351 S-Timer Antigens in Mice. Humoral immune responses in Stage 2 of the study were evaluated on Day 49 (2 weeks after dose 3) based on SARS-CoV-2 pseudovirus neutralization assays against (**a**) Original (Wuhan-Hu-1) strain, (**b**) UK (B.1.1.7) variant, (**c**) South African (B.1.351) variant and (**d**) Brazil (P.1) variant pseudoviruses. Results here are shown for all subgroups (representing all vaccination combinations) in the study (n=4/subgroup). Dots represent individual animals; bars represent geometric mean titers (GMT) of EC_50_ values, and error bars represent 95% confidence intervals (95% CI).

**Table S1.**
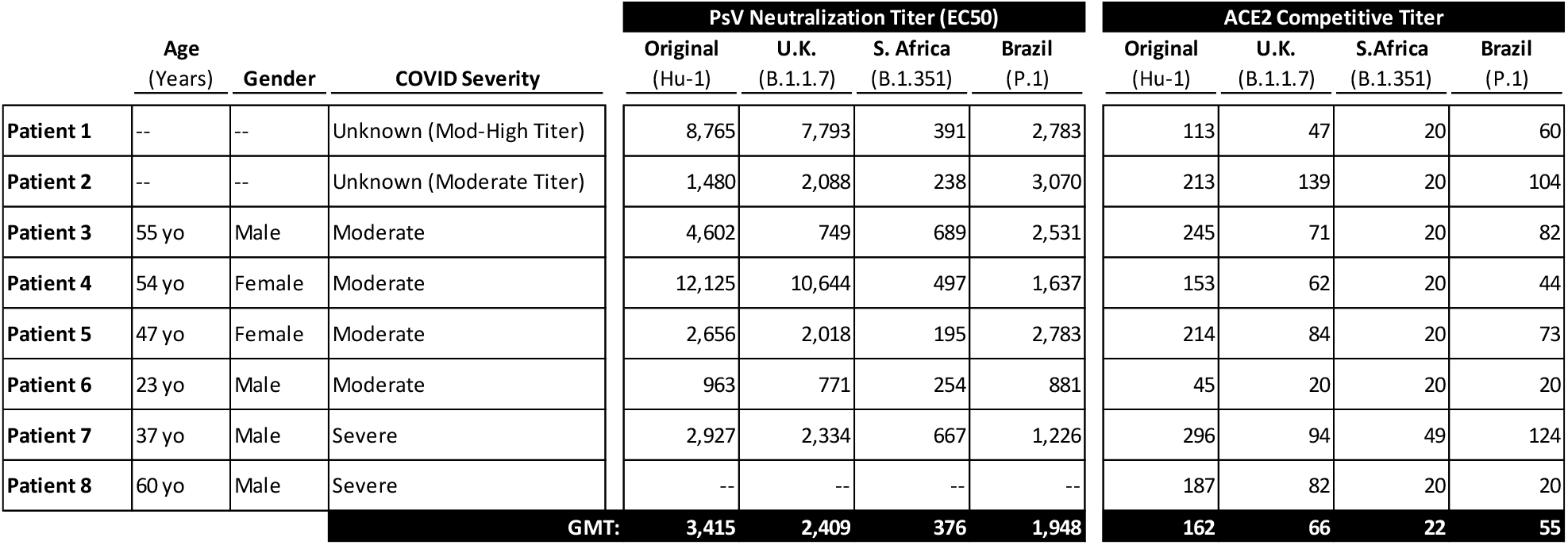
Human convalescent sera panel. Characteristics and information of 8 COVID-19 patients from whom convalescent sera was collected and included for testing in this study. Patient age, patient gender and COVID-19 disease severity (if available) are included, as well as SARS-CoV-2 pseudovirus neutralization titers (EC_50_) and ACE2-competitive titers (EC_50_) against the original strain, UK variant (B.1.1.7), South African variant (B.1.351) and Brazil variant (P.1) strains.

